# Environmental Persistence and Disinfection of Lassa Virus

**DOI:** 10.1101/2023.05.17.541161

**Authors:** Marlee Shaffer, Robert J. Fischer, Shane Gallogly, Olivia Ginn, Vincent Munster, Kyle Bibby

## Abstract

Lassa Fever, caused by Lassa virus (LASV), is endemic to West Africa, where approximately 300,000 illnesses and 5,000 deaths occur annually. LASV is primarily spread by infected multimammate rats via urine and fomites, highlighting the importance of understanding the environmental fate of LASV. This study evaluated the persistence of LASV strains on surfaces in aqueous solutions and with sodium hypochlorite disinfection. LASV strains (Josiah and Sauerwald) were more stable in DI water (k = 0.23 and 0.34 days^-1^) than primary influent wastewater (k = 1.3 and 1.9 days^-1^). The decay rates of LASV on HDPE (k = 4.3 and 2.3 days^-1^) and Stainless Steel (k = 5.3 and 2.7 days^-1^) were not significantly different for either strain. Sodium hypochlorite was highly effective at inactivating both strains of LASV. This work presents data for the environmental persistence of LASV to inform future risk assessment and management efforts.

## Introduction

Lassa Fever is an acute viral hemorrhagic illness caused by Lassa virus (LASV), an enveloped RNA virus of the *Arenaviridae* family. (1) Lassa Fever is endemic to West Africa, resulting in 300,000 cases and 5,000 deaths annually. (1,2) Over the past decade, there has been an increase in Lassa Fever cases, likely attributed to an increasing population, urbanization, and environmental changes. (3) Approximately 80% of Lassa Fever cases result in mild symptoms; however, the remaining 20% are serious infections causing hemorrhaging and vomiting while simultaneously impacting several organs, including the kidneys, liver, and spleen, which can have lifelong effects for survivors. (4) The overall mortality rate for Lassa Fever is 1%; however, patients hospitalized with a severe infection have an increased mortality rate of 15 to 20%. (1,5)

The natural reservoir of LASV is the multimammate rat, *Mastomys natalensis*, found in 50 to 98% of households in West Africa, putting approximately 58 million people at risk of infection. (2,4,6) Ecological niche models predicting expansion of suitable environmental conditions for LASV coupled with population increases in these areas estimate a 761% increase in the risk of infection for people in endemic areas. (7) Transmission of LASV is primarily through direct contact with fomites contaminated with rat urine or feces or inhaling fomites. (2,8) Secondary person-to-person transmission can occur from direct contact with bodily fluids of infected individuals, specifically in healthcare settings with inadequate infection prevention and control measures. (9) Thus, understanding the environmental persistence of LASV is critical for mitigation and control efforts.

The World Health Organization lists LASV as a priority pathogen with pandemic potential, as there is a relatively long incubation period and a lack of prior immunity within populations outside of endemic areas. (10) Infected individuals shed the virus in bodily fluids for up to three months. (10) There is no standard practice for disposing of LASV infectious waste, leading to concerns about further environmental transmission. Limited research is available evaluating the environmental persistence and disinfection of infectious LASV, creating a critical gap in understanding the potential for continued environmental transmission. Here, we investigated the persistence of two LASV strains on representative surfaces and in water and wastewater. We then assessed LASV disinfection with sodium hypochlorite to assess a potential LASV management approach.

## Materials and Methods

### Biosafety Statement

The experiments were conducted in a BSL4 facility, and the samples were removed following approved standard operating procedures.

### Surface Persistence Experiments

Solid surface persistence experiments were conducted similarly as previously described. (15) Briefly, 3 disks (4 cm diameter) of each material were placed individually into wells of a 6-well plate for each time point. 10^6^ TCID_50_ of either Josiah (lineage IV) or Sauerwald (lineage II) LASV isolates in the cell-free medium were evenly distributed on the disks, and the plates were allowed to air dry at 20 °C. At each time point, 450 μL of DMEM (Sigma) modified as described above were placed onto the surface and gently agitated to free the surface-bound virus. Viral titers for each surface were determined as described below.

### Aqueous Persistence Experiments

All experiments were completed in triplicate and conducted at 20 °C. Josiah and Sauerwald LASV stocks were added to each replicate of DI water and wastewater to achieve a final concentration of 10^6^ TCID_50_ mL^-1^. Samples were collected daily for five days. At each time point, including the time zero, 50 μL of the aqueous matrix from each of the bulk wastewater or DI water vials was added into 450 μL of Dulbecco’s modified Eagle’s medium (DMEM, Sigma) supplemented with heat-inactivated fetal bovine serum (FBS, Gibco) and 2%, Pen/Strep (Gibco) to a final concentration of 50 U/mL penicillin and 50 μg/mL streptomycin, and L-glutamine (Gibco) to a final concentration of 2 mM in an appropriately labeled 2 mL screw top vial and frozen at -80 °C. Negative controls were 50 μL of the non-spiked aqueous matrix. Viral titers for each replicate were determined as described below.

### Aqueous Disinfection Experiments

The initial chlorine demand of the wastewater matrix was assessed using the Hach Free Chlorine test kit (Method 10069). (11) LASV disinfection was assessed, in triplicate, in wastewater for two isolates, Josiah and Sauerwald, at 1 mg/L, 5 mg/L, and 10 mg/L, of sodium hypochlorite. In the top row of a deep-well 96-well plate virus was spiked into wastewater or DI water to an initial concentration of 10^6^. An initial sample was removed, diluted 1:10 in DMEM and frozen at -80°C. Sodium hypochlorite (2 μL) was then added to each well to achieve the desired NaO_4_Cl concentrations. The ‘time zero’ sampling point was taken approximately 20 seconds following the addition of chlorine to enable sample mixing. Samples (50 μL) were removed at 1, 5, 15, and 30 minutes, mixed with 50 μL of sodium thiosulfate, concentration matched (w:w) with the sodium hypochlorite, to quench any remaining free chlorine. This was then added to 400 μL of DMEM to achieve a final 1:10 dilution prior to freezing at -80°C until titrated.

### Virus Titration

To assess the concentration of viable virus in each sample a tissue culture infectious dose 50 (TCID_50_) was performed using 4 10X dilution series for each. Vero E6 cells were then incubated for 1 hour with the virus dilutions at 37°C at which point the virus was removed from the two highest concentrations, rinsed two times with PBS, and 200 μL of fresh culture medium was added. Fresh culture medium (100 μL) was also added to the remaining wells in the plate. The plates were incubated at 37 °C for seven days, inspected for cytopathic effect (CPE), and scored using the method of Spearman-Kärber.

### Statistical Analysis

The decay of infectious LASV strains for all experiments was analyzed using a first-order decay model excluding data points below the limit of detection for the assay, as done previously. (11–14) Each plotted point represents the mean of three experimental replicates, and the error bars show the standard deviation. The equation of the linear regression and the R-squared value are shown on the plots. All plotting, regressions, and statistical analyses were performed using R Studio 2022.07.2.

## Results

### Persistence of Lassa Virus on Surfaces

Stainless steel and high-density polyethylene (HDPE) are commonly used non-porous materials for hospital equipment, additionally, HDPE is a component of common full-body personal protective coveralls and gowns worn in hospitals. (19) LASV was deposited on HDPE and stainless steel surfaces, allowed to air dry at 20 °C, and virus viability was measured over five days. **Figure 1** shows the first-order decay of the Josiah and Sauerwald isolates on HDPE and stainless steel. No significant difference in decay was found between the HDPE and stainless steel within a single strain. Conversely, there was a small but significant difference between the two strains on Stainless Steel (p = 0.028) but not on HDPE.

**Table 1.**
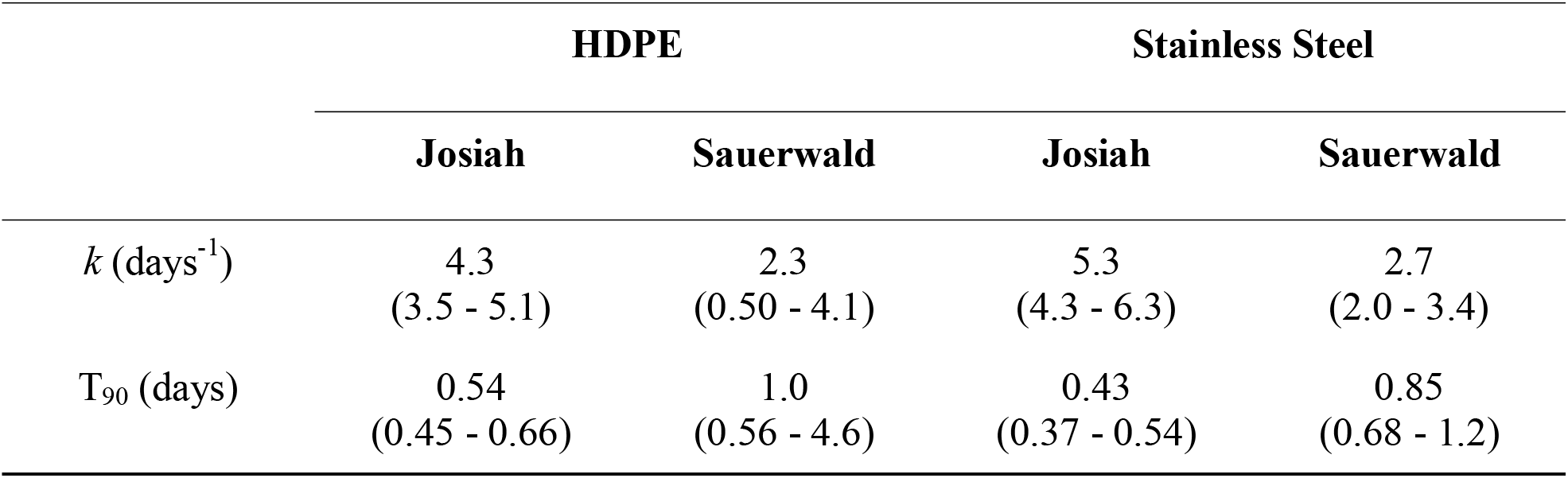
First-Order Decay Rate Constants and Decimal Reductions with 95% Confidence Intervals for LASV Strains on Surfaces

|  | HDPE |  | Stainless Steel |  |
| --- | --- | --- | --- | --- |
|  | Josiah | Sauerwald | Josiah | Sauerwald |
| $k$ (days<sup>-1</sup>) | 4.3<br>(3.5 - 5.1) | 2.3<br>(0.50 - 4.1) | 5.3<br>(4.3 - 6.3) | 2.7<br>(2.0 - 3.4) |
| $T_{90}$ (days) | 0.54<br>(0.45 - 0.66) | 1.0<br>(0.56 - 4.6) | 0.43<br>(0.37 - 0.54) | 0.85<br>(0.68 - 1.2) |

**Figure 1.**
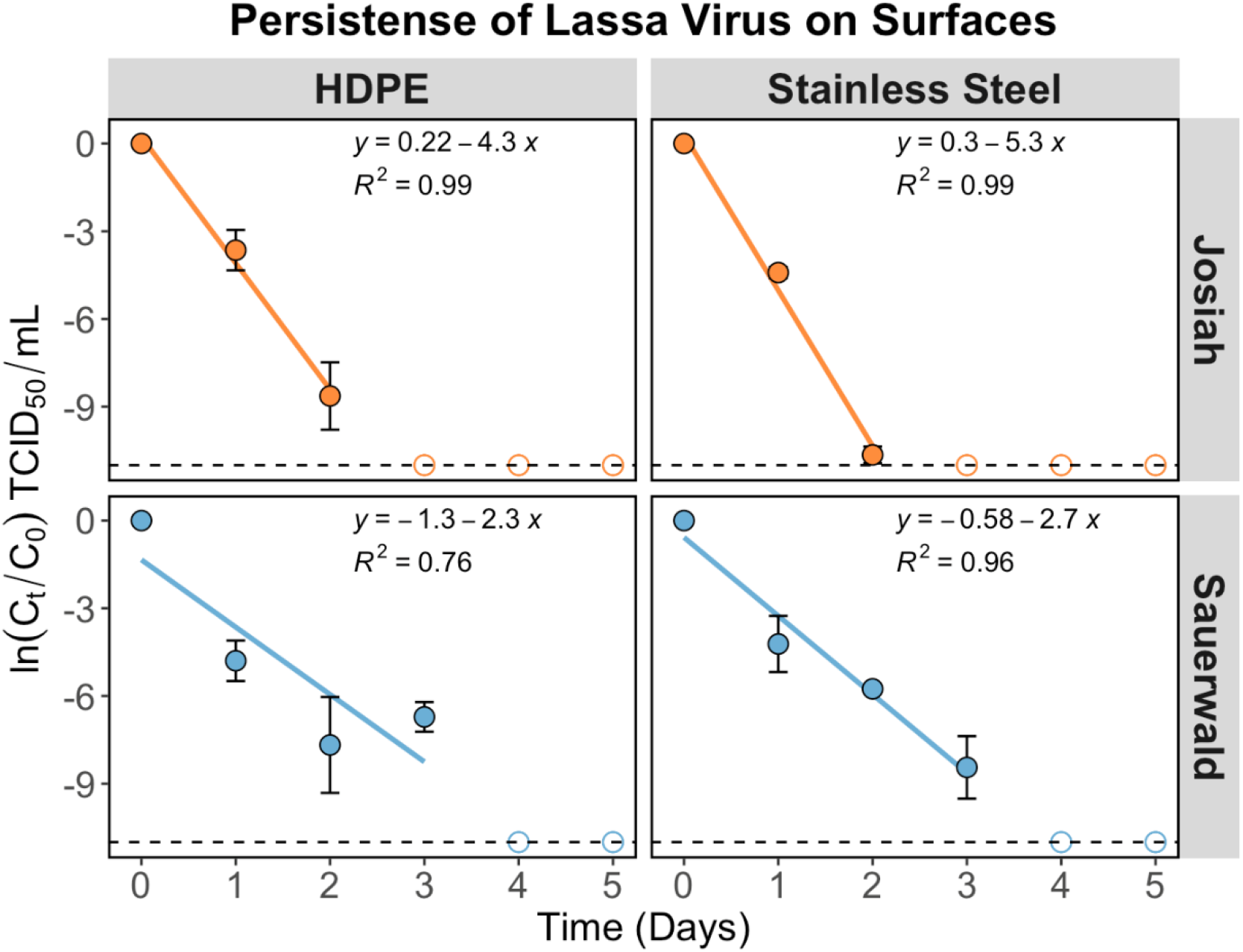
First-order decay of Josiah and Sauerwald LASV isolates on HDPE and stainless steel. The x-axis shows the time in days, and the y-axis shows the normalized natural log of the observed TCID_50_ per mL decay. The dashed line shows the assay’s detection limit, and hollow circles show points below the detection limit, which were not included in the regression analysis.

### Persistence of Aqueous Lassa Virus

LASV strains were spiked into DI water and raw municipal wastewater, and persistence was monitored for five days. **Figure 2** shows the first-order decay for each strain and condition. For both strains, the decay rate was significantly higher in wastewater than in DI water (p < 0.002), which can be attributed to multiple factors. (15) First, microorganisms found in wastewater contribute to viral inactivation. (16) Second, RNA genomes experience increased degradation when exposed to higher concentrations of ammonia found in wastewater. (17) No significant difference existed between LASV strains for DI water or wastewater. **Table 2** summarizes the first-order decay rates and the decimal reduction, T_90_, for LASV in DI water and wastewater.

**Table 2.** First-Order Decay Rate Constants and Decimal Reductions with 95% Confidence Intervals for Aqueous LASV Strains

|  | DI Water |  | Wastewater |  |
| --- | --- | --- | --- | --- |
|  | Josiah | Sauerwald | Josiah | Sauerwald |
| $k$ (days<sup>-1</sup>) | 0.23<br>(0.04 - 0.43) | 0.34<br>(0.15 - 0.53) | 1.3<br>(0.94 - 1.6) | 1.9<br>(1.5 - 2.3) |
| $T_{90}$ (days) | 10.0<br>(5.4 - 57.6) | 6.8<br>(4.3 - 15.4) | 1.8<br>(1.4 - 2.4) | 1.2<br>(1.0 - 1.5) |

**Figure 2.**
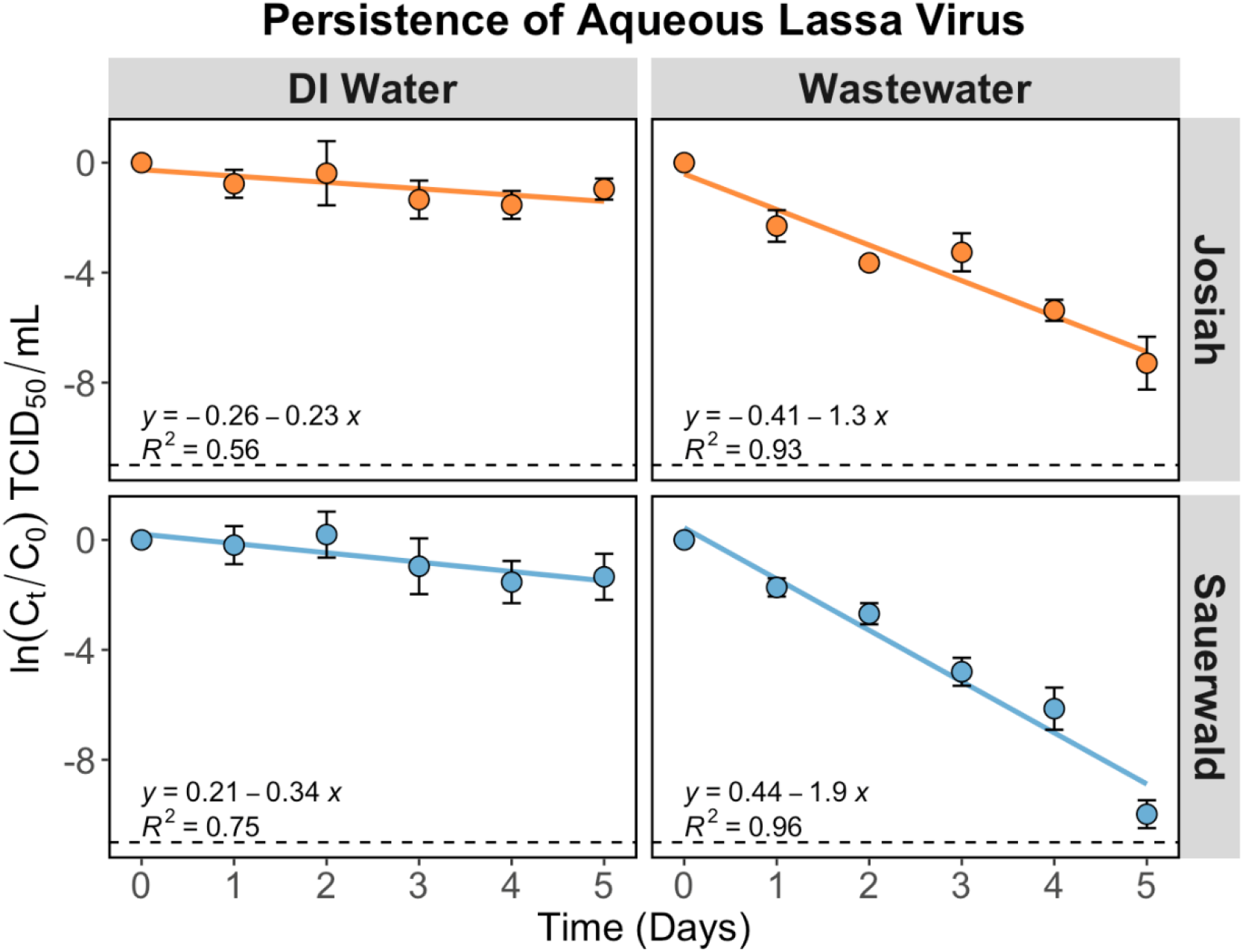
First-order decay of Josiah and Sauerwald strains in DI water and wastewater. The x-axis shows the time in days, and the y-axis shows the natural log of the observed TCID_50_ per mL decay. The dashed line shows the limit of detection of the assay.

### Disinfection of Lassa Virus with Sodium Hypochlorite

Sodium hypochlorite is a widely available disinfectant, effective at inactivating microorganisms, and produces a disinfectant residual resulting in its widespread use for wastewater disinfection. (18) Hospitals have also used sodium hypochlorite to manage wastewater containing high-consequence pathogens on-site. (19) Prior LASV disinfection studies have evaluated thermal inactivation, UV irradiation, and gamma irradiation in human serum and plasma. (20) The disinfection kinetics of Josiah and Sauerwald LASV strains were assessed in raw municipal wastewater at 0 mg/L, 1 mg/L, 5 mg/L, and 10 mg/L sodium hypochlorite (**Figure 3**).

**Figure 3.**
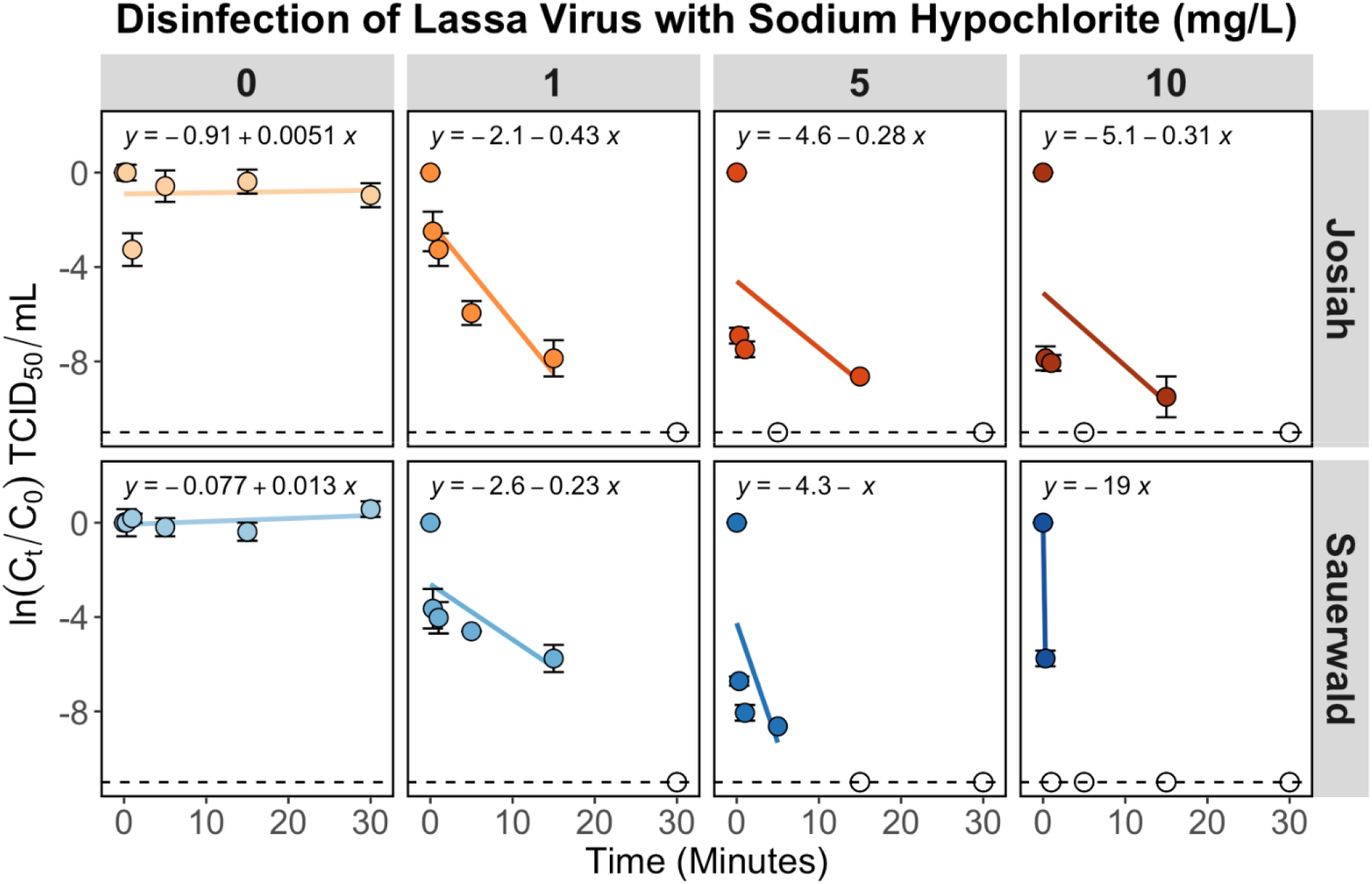
Disinfection of Lassa virus with sodium hypochlorite in municipal wastewater. The x-axis shows the time in days, and the y-axis shows the normalized natural log of the observed TCID_50_ per mL decay. The dashed line shows the assay’s detection limit, and hollow circle represent where all data points were below the detection limit, which were not included in th regression analysis.

There was no removal at 0 mg/L sodium hypochlorite over the experimental timeframe for either strain. Increases in sodium hypochlorite concentrations resulted in significantly higher decay rates for Josiah (p = 0.001 - 0.3) and Sauerwald (p = 0.001 - 0.007) strains of LASV. The Sauerwald strain showed faster inactivation than the Josiah strain for both 5 mg/L and 10 mg/L (p = 0.04). These results indicate that sodium hypochlorite effectively inactivates the LASV strains analyzed in this study.

## Discussion

Generally, LASV persistence was longer in aqueous solution than on surfaces. The persistence of the Josiah strain was significantly longer in wastewater than on HDPE and stainless steel (*p =* <0.01). Both Josiah and Sauerwald strains had significantly lower decay rates in DI water compared to the decay on HDPE and stainless steel (*p =* < 0.01 - 0.02), indicating higher stability in DI water than on both surfaces.

Contact with potentially contaminated surfaces without proper personal protective equipment, especially in hospitals, is anticipated to be a potential transmission route for LASV (21); however, no previous studies have analyzed the persistence of infectious LASV on surfaces. Quantifying LASV persistence is critical to implementing proper control mechanisms and prevention measures. (22) The persistence of enveloped viruses on surfaces is specific to the virus, the surface, and the temperature and humidity used. A previous study using Ebola virus at 21°C and 40% RH on HDPE and Stainless Steel found the decay of Ebola to be slower than we found for LASV on the same surfaces. (12) In contrast, a study by van Doremalen et al. found the decay rates of SARS-CoV-1 and SARS-CoV-2 on Stainless Steel to be faster than the Sauerwald strain but slower than the Josiah strain of LASV from this study. (23) Together with the increased LASV persistence in aqueous solution compared to surfaces, this data suggests that efforts are best directed at managing fluids containing infectious LASV recently deposited onto surfaces.

Bacteriophage Phi6 and Murine Hepatitis Virus are commonly used as surrogates for enveloped viruses of concern, such as Ebola and SARS-CoV-2, and potentially LASV. Bacteriophage Phi6 and Murine Hepatitis Virus decay rates were previously assessed in wastewater. Compared to the LASV decay rates from this study, surrogate decay was 2 – 5 times faster than LASV, indicating surrogate viruses are not representative of LASV persistence in wastewater. (24) Furthermore, in DI water, the decay rate for bacteriophage Phi6 was faster than that found for LASV, showing lower surrogate virus persistence. (15) Conversely, in autoclaved wastewater influent, the decay rate of bacteriophage Phi6 was lower than what we found for LASV strains, suggesting potential differences in the decay of viruses based on the specific matrix. (15) Overall, commonly used surrogates for enveloped viruses of concern do not indicate the behavior of LASV in aqueous solutions.

Wastewater in LASV endemic locations will likely be of higher strength (i.e., more concentrated) than the primary influent wastewater used here. Wastewater from a LASV treatment unit will present higher ammonia levels, total suspended solids, and biological oxygen demand. These physiochemical changes would be expected to decrease LASV persistence but increase the amount of sodium hypochlorite needed to reach similar free chlorine levels due to increased chlorine demand. If sodium hypochlorite is used for disinfection, then chlorine residual should be assessed to ensure proper sodium hypochlorite concentrations for LASV inactivation. In addition, the two strains evaluated displayed strain-dependent behavior suggesting that the potential for environmental transmission may vary by strain. Future studies should investigate the mechanistic basis of this differing strain behavior.

Our findings supply data for the environmental persistence of LASV, a WHO priority pathogen with pandemic potential, on representative surfaces, in aqueous solutions, and with sodium hypochlorite disinfection. LASV showed shorter persistence on surfaces than previously assessed enveloped virus surrogates and priority pathogens, and longer persistence in aqueous solution. LASV was rapidly disinfected in the presence of free chlorine from the addition of sodium hypochlorite. This study provides critical data about the stability and control of LASV in the environment to further management and mitigation efforts.

## Acknowledgements

This work was supported by the Intramural Research Program of the National Institute of Allergy and Infectious Diseases (NIAID), National Institutes of Health (NIH) and Water Research Foundation award 5029.

