## Supporting Information for "Environmental Persistence and Disinfection of Lassa Virus"

**Table S1:** Composition of Tested Wastewater

**Figure S1:** Free Chlorine Concentration

*Wastewater Characterization*

Approximately 1L of primary influent wastewater was aseptically collected from a wastewater treatment plant in northern Indiana, United States, and stored at -80°C before being shipped overnight on ice to Rocky Mountain Laboratories (RML). All physiochemical characterization and chlorine demand testing were conducted at the University of Notre Dame. **Table S1** describes the physiochemical characteristics of the influent wastewater used in the following experiments.

**Table S1.** Composition of Tested Wastewater

| Constituent | Tested Wastewater |
| --- | --- |
| pH | 6.16 |
| Chemical Oxygen Demand (mg/L) | 279 |
| Ammonia (mg/L) | 29 |
| Nitrate (mg/L) | 5.0 |
| Phosphorus (mg/L) | 9.2 |
| Total Suspended Solids (mg/L) | 134 |

*Chlorine Demand and Ct Values*

The initial free chlorine residuals in wastewater were dose-dependent, with the 0, 1, 5, and 10 mg/L doses resulting in initial free chlorine residuals of 0, 0.97, 3.43, and 6.53 mg/L, respectively. The free chlorine measurements are shown in **Figure S1**.

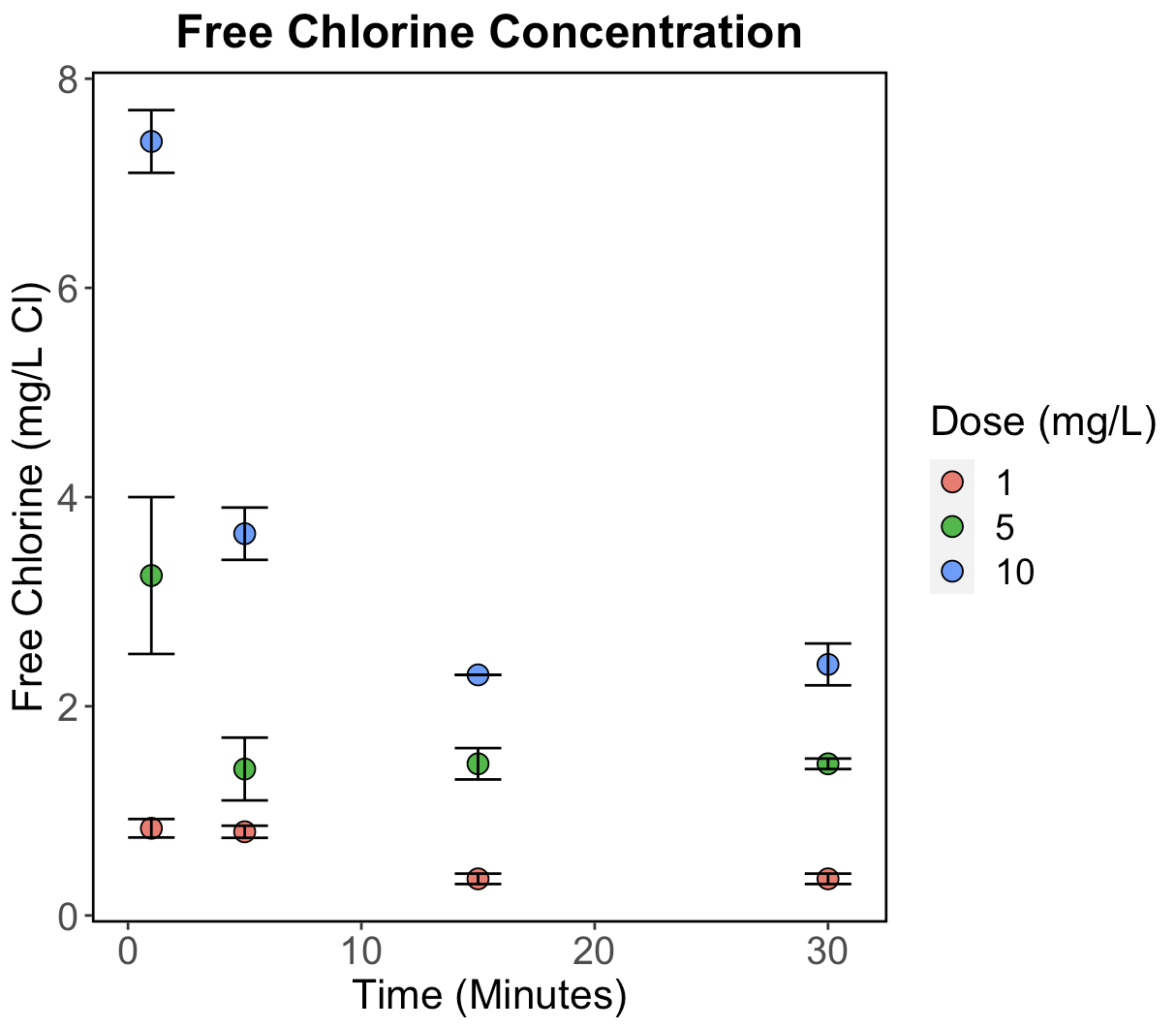

**Figure S1.** Free Chlorine Concentrations for 1, 5, and 10 mg/L of free chlorine as sodium hypochlorite. The y-axis shows the free chlorine concentration as mg/L of Cl determined by the HACH D900 Colorimeter. The x-axis shows the time in minutes. Each data point represents the mean of three replicates, and the error bars show the standard deviation.
